# Clinical significance of C4d deposition in renal tissues from patients with primary Sjögren’s syndrome—A preliminary study

**DOI:** 10.1101/562215

**Authors:** Wenli Xia, Bixia Gao, Lin Duan, Yan Li, Yubing Wen, Limeng Chen, Xuemei Li, Falei Zheng, Mingxi Li

**Author notes:** Corresponding author: Mingxi Li, No. 1 Shuaifuyuan, Wangfujing Street, Beijing, 100730, China.

## Abstract

**Objectives:** To evaluate renal expression of C4d, a complement component in the classical/mannose binding lectin (MBL) pathway, in patients with primary Sjögren’s syndrome (pSS)-associated renal impairments.

**Methods:** We retrospectively reviewed the clinical and pathological data from 39 patients with pSS presenting with renal impairments. C4d was examined in paraffin-embedded biopsy tissues using immunohistochemistry. Glomerular C4d positive was defined when >75% glomeruli were globally stained. Tubulointerstitial C4d (TI-C4d) were scored semi-quantitatively as 0 (absent), 1 (spotty or weak), 2 (patchy) and 3 (diffuse). A TI-C4d score ≥2 was considered TI-C4d positive and included in the TI-C4d^+^ group and vice versa. Peritubular capillary (PTC) C4d was scored as 0 (absent), 1 (0∼10%, minimal), 2 (10%∼50%, focal), and 3 (>50%, diffuse).

**Results:** Glomerular C4d deposition was observed in all 8 patients with pSS-related membranous nephropathy (MN) without obvious C1q deposition. Two of 5 patients with mesangial proliferative glomerulonephritis and 1 of 2 patients with IgA nephropathy had mild mesangial C4d deposition. Sixteen patients (6 glomerular dominant and 10 tubulointerstitial dominant) presented TI-C4d score ≥2. Patients in the TI-C4d^+^ group exhibited a higher serum creatinine level at the time of renal biopsy (TI-C4d^+^ 132.5 [89.7, 165.5] vs. TI-C4d^-^ 83.0 [70.7, 102.0] μmol/L, P=0.008). PTC C4d was observed in 12 patients, with each of minimal, focal and diffuse staining being noted in 4 patients.

**Conclusions:** The MBL pathway of complement activation was potentially involved in pSS-related MN. Tubulointerstitial C4d might be a pathological marker of severe renal injury in patients with pSS-related renal impairments.

## Background

The kidney is affected in approximately 0.3% to 33.5% of patients with primary Sjögren’s syndrome (pSS), a systemic disease characterized by sicca symptoms and anti-Ro/SSA and/or anti-La/SSB antibodies(Francois &Mariette, 2016;Garcia-Carrasco et al., 2002;Ramos-Casals et al., 2014). Two main pathological entities have been described. Tubular interstitial nephritis (TIN) is the major type and is often characterized by renal tubular acidosis (RTA) with or without renal insufficiency(Evans et al., 2016). Glomerulonephritis (GMN) is the other type, including membranoproliferative glomerulonephritis (MPGN), membranous nephropathy (MN), mesangial proliferative glomerulonephritis (MePGN), focal segmental glomerular sclerosis (FSGS) and minimal change disease (MCD)(Jasiek et al., 2017;Kidder et al., 2015;Maripuri et al., 2009). MN (pSS-MN) is one of the major types of GMN reported in different studies (Carrillo-Pérez et al., 2018;Francois &Mariette, 2016).

Primary membranous nephropathy (PMN) is a kidney-specific disease with increased proteinuria and pathological characteristics of sub-epithelial immune complex deposition (predominantly IgG4 and C3 deposition)(Sinico et al., 2016). PMN is characterized by the presence of serum autoantibodies against the podocyte component M-type phospholipase A2 receptor (PLA2R) in approximately 70% of patients(Beck et al., 2009) and anti-thrombospondin type 1 domain containing 7A (THSD7A) antibodies in approximately 10% of those with negative anti-PLA2R antibodies (nearly 3% of patients with intrinsic antibodies related MN)(Hayashi et al., 2018;Tomas et al., 2014). The mannose-binding lectin pathway of complement activation is involved in PMN, as evidenced by sub-epithelial deposition of hypogalactosylated IgG4 and C3 and 100% glomerular C4d deposition without C1q co-deposition(Hui et al., 2014).

Notably, pSS is one of the possible causes of secondary MN(Francois &Mariette, 2016). Patients with pSS-MN may present extra renal manifestations that do not exist in patients with PMN(Goules et al., 2013;Maripuri et al., 2009). In patients with MN secondary to pSS, the complement activation pathway has not been studied. Patients with pSS-TIN are characterized by interstitial lymphocyte infiltration with rare tubular and interstitial immune complex deposition(Bossini et al., 2001). Autoantibodies against tubular components have been identified in patients with pSS-TIN by some authors(Devuyst et al., 2009;Takemoto et al., 2005;Takemoto et al., 2007), and the complement component C9 was also observed around TBM in some patients with pSS-TIN(Evans et al., 2016). It has not extensively researched whether immune complex-mediated interstitial injury plays a role in pSS-TIN.

C4d deposition in various compartments of renal tissues have been reported. C4d deposition in the glomerulus indicates local complement activation through the classical or lectin pathway(Ehrnthaller et al., 2011). It is used to differentiate C3 glomerulopathy and immune complex-mediated proliferative GN(Sethi et al., 2015). Mesangial C4d deposits have been observed in patients with IgA nephropathy (IgAN)(Espinosa et al., 2009), lupus nephritis (LN)(Kim et al., 2013), and thrombotic microangiopathy (TMA)(Chua et al., 2015). Peritubular capillary (PTC) C4d correlates with antibody-mediated transplant rejection and inferior renal allograft outcomes(Regele et al., 2002). Tubular C4d deposition is associated with an increased WHO grade of IgAN(Maeng et al., 2013). To our knowledge, neither glomerular C4d deposition nor tubulointerstitial C4d deposition have been investigated in patients with pSS-MN and in patients with pSS-TIN.

In the present study, we performed a retrospective examination of 39 patients with pSS with renal involvement (21 cases with TIN and 18 cases with GMN) from a single centre. C4d were examined in formaldehyde-fixed, paraffin-embedded biopsy tissues using immunohistochemistry. The objectives were to investigate the prevalence and localization of C4d deposits in renal biopsy tissues in patients with pSS-MN and those with pSS-TIN.

## Methods

### 1. Patient selection

We retrospectively analysed the clinical and pathological data from 82 hospitalized patients with biopsy-proven renal involvements secondary to “Sjögren’s syndrome” (ICD: M35.001) in Peking Union Medical College Hospital (PUMCH) from 1996 to 2011. After Sjögren’s syndrome was redefined based on the 2002 American and European Classification Criteria (AE), we excluded 13 patients with systemic lupus nephritis (LN), 2 patients with primary biliary cirrhosis (PBC), 3 patients with rheumatoid arthritis (RA), 2 patients who were positive for anti-HCV antibodies, 2 patients with unidentified GMN due to glomerular sclerosis, and 3 patients for whom adequate kidney biopsy samples were unavailable. In addition, 18 patients who only qualified based on the Fox criteria which does not require the presence of anti-SSA/SSB antibodies were excluded. Thirty-nine patients were enrolled (Fig 1). All inpatients received a consultation with a rheumatologist and had a confirmed diagnosis of primary Sjögren’s syndrome. After patient selection, archived formalin-fixed and paraffin-embedded kidney tissues were retrieved and stained as described below. The study was performed in accordance with the ethical standards of the Declaration of Helsinki and was approved by the Institutional Review Board of PUMCH (Reference code:S-k585).

**Figure 1.**
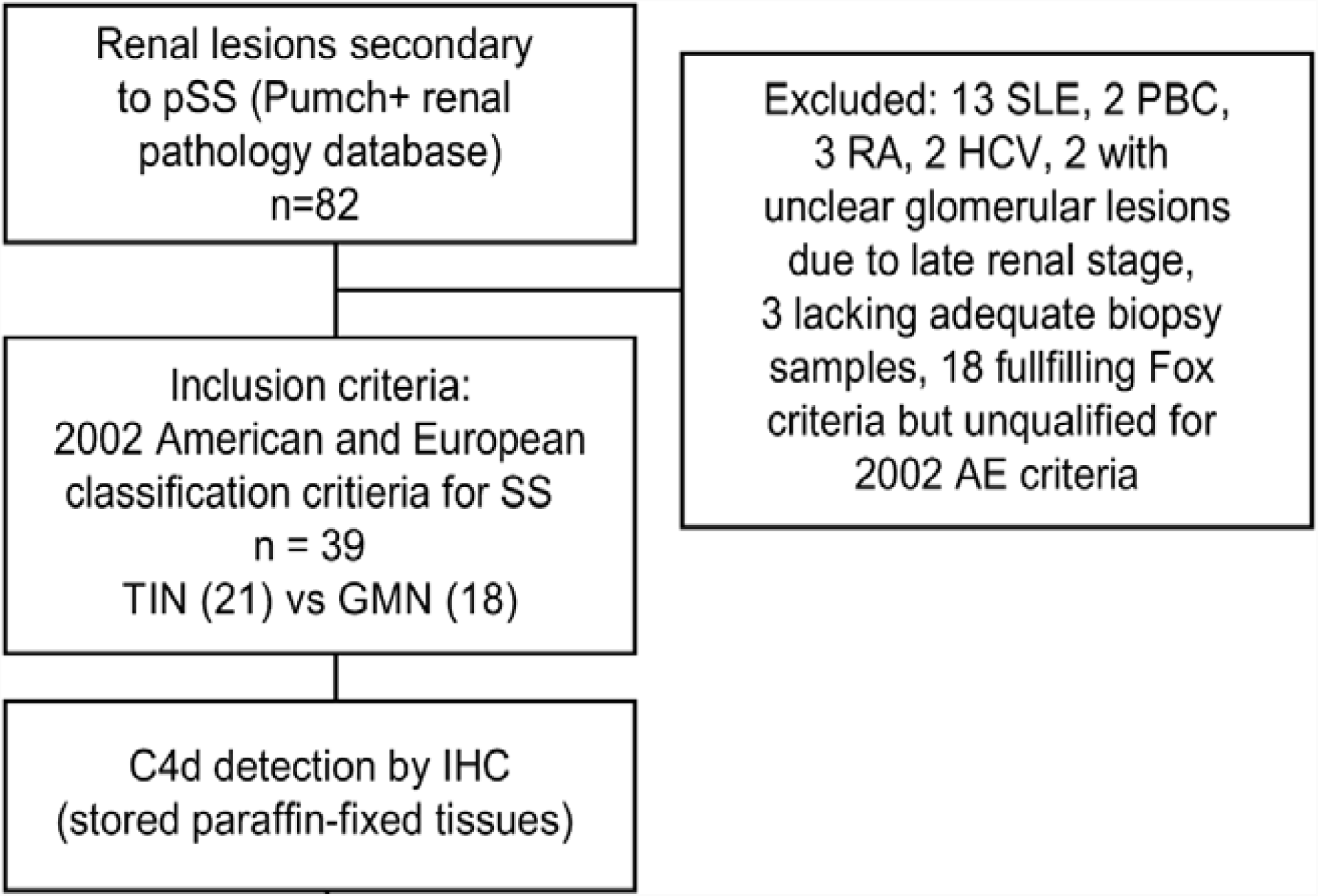
Patients selection chart.

### 2. Clinical data

Medical records including the demographic data, the duration of pSS, the duration of renal impairment from symptom onset to renal biopsy, medical comorbidities and treatment history were retrospectively reviewed. The immunological screening for antinuclear antibodies (ANA), anti-SSA/SSB antibodies, immunoglobulins (IgG, IgM and IgA), complement (CH50, C3 and C4), C-reactive protein (CRP), rheumatoid factor (RF) and cryoglobulins were obtained.

Renal function was evaluated by the estimated glomerular filtration rate (eGFR) with the CKD Epidemiology Collaboration (CKD-EPI) equation. Renal tubular function screening included blood and urine pH values, electrolytes, blood and urine α1-macroglobulin and β2-microglobulin level, urine transferrin protein and N-acetyl-β-amino-glucosidase levels. Ammonium chloride load test and sodium hydrogen carbonate reabsorption rate were applied to verify the diagnosis of renal tubular acidosis. Urine and blood osmolality, fluid deprivation/vasopressin test and cranial images were evaluated to assess the renal tubular concentration function and to differentiate diabetes insipidus. Proteinuria based on 24-hour urine protein excretion rate was classified as mild (<1.5 g/day), moderate (1.5-3.5 g/day) and nephrotic (>3.5 g/day). Urinary ultrasound results were reviewed for evidence of anomalies and renal calcification.

### 3. Kidney pathology and C4d detection

#### (1) Pathological examination of kidney tissue

The renal histological diagnosis was based on light microscopy (LM), immunofluorescence (IF) staining and electron microscopy (EM) findings. The diagnosis was made by two independent pathologists. Patients were divided into the TIN group (with TIN only) and GMN group (including patients with interstitial lesions) based on pathological findings.

Kidney biopsy specimens were scored semi-quantitatively: (1) mesangial proliferation: 0 (no proliferation, ≤3 mesangial cells per mesangial area), 1 (mild to moderate proliferation, 4-6 mesangial cells per area) and 2 (intense proliferation, >6 mesangial cells per area). (2) Glomerular sclerosis and crescent were counted as a percentage. Eight cases with no more than 15 glomeruli per specimen were only excluded from the calculation of this item. (3) Tubular atrophy and interstitial fibrosis were scored according to the percent area involved: 0 (absent), 1 (≤25%), 2 (25-50%), 3 (50%-75%), and 4 (>75%). IgG, IgM, IgA, C1q, C3, C4, fibrinogen, and light chain κ and λ levels were detected using direct IF staining and scored as 0 (negative), 1 (1+), 2 (2+), 3 (3+) and 4 (4+). IF-positive samples were defined as positive for IgG, IgM, IgA, C1q, C3, C4, κ or λ and scored as ≥1 point.

#### (2) Immunohistochemical staining of C4d

Rabbit anti-human polyclonal C4d antibodies (1:40; Biomedica, Vienna, Austria) were applied on 2-μm formaldehyde-fixed kidney sections. Antigen retrieval was performed by incubating the sections with 0.4% pepsin (Zhongshan Golden Bridge Biotechnology, Beijing, China) for 30 min at 37°C. Sections were incubated with the primary antibody overnight at 4°C and with avidin-free horseradish peroxidase (HRP)-conjugated goat anti-rabbit immunoglobulins using the two-step Envision kit (PV9001, Zhongshan Golden Bridge Biotechnology, Beijing, China) for 30 min at 37°C. Sections were treated with a freshly prepared 3-3-diaminobenzidine solution (DAB) (Zhongshan Golden Bridge Biotechnology, Beijing, China) for 3 min and counter-stained with haematoxylin for 30 seconds.

A positive control for C4d was obtained from dendritic lymphocytes in human tonsil tissues from a patient undergoing an operation for obstructive sleep apnoea hypopnea syndrome(Fig S1)(Zwirner et al., 1989). Renal tissues from patients with mild lesions who underwent a biopsy for microscopic haematuria were used as negative controls (Fig S2).

#### (3) C4d evaluation and scoring in different compartments

Glomerular C4d positive (G-C4d^**+**^) was defined when >75% glomeruli had global staining (involving >50% area of one glomerulus),vice versa glomerular negative (G-C4d^**-**^)(Espinosa et al., 2009).

Tubulointerstitial C4d (TI-C4d) deposition was semi-quantitatively scored as 0 (absent), 1 (weak or spotty stain), 2 (patchy stain), and 3 (diffuse stain) within microscopic fields. A TI-C4d score >1 was defined as positive (TI-C4d^+^)(Tan et al., 2011). Using a TI-C4d deposit score of >1 as a cut-off, patients were classified into the TI-C4d^+^ TI-C4d^**-**^ groups.

Peritubular capillary C4d (PTC-C4d) staining was evaluated and graded according to the Banff criteria as negative (0, absent), minimal (1, less than 10%), focal (2, between 10% and 50%) and diffuse (3, >50%)(Haas et al., 2014).

### 4. Statistics

Continuous variables with a normal distribution were analysed using Student’s t-test or the Mann-Whitney U-test for abnormally distributed data. Categorical data were compared using the Fisher’s exact test. Correlations were analysed by calculating Pearson’s correlation coefficients for continuous normally distributed variables, Spearman’s Rho for nominal and ordinal variables, and performing a linear regression analysis for continuous variables. P<0.05 was considered statistically significant. Calculations were performed using SPSS statistical software version 11.5 (SPSS, Chicago, IL, USA).

## Results

### 1. Demographic characteristics of patients with pSS presenting renal involvement

Among the 39 patients included in this study, 21 were diagnosed with TIN, and 18 were diagnosed with GMN. Eight of 18 patients with GMN also had TIN. The male:female ratio was 9:30. The average age of the patients was 42±14.1 (9-72) years at the time of renal biopsy. ANA was positive in 37/39 patients, with different titres: 8 patients at a 1:1280 titre, 8 patients at 1:640, 9 patients at 1:320, 9 patients at 1:160, 2 patients at 1:80, and 1 patient at 1:40. Hyper-γ-globulinemia was present in 11/36 (28.2%) patients. Elevated serum Ig levels and hypocomplementemia were also observed (Table 1). Twenty-two (56.4%) patients were diagnosed with RTA (18 type I cases and 4 type II cases). Three of the 4 patients with type II RTA simultaneously exhibited Fanconi syndrome. Twenty (48.8%) patients had polynocturia, and two had diabetes insipidus.

**Table 1:** Demographic, clinical and serological characteristics of patients with pSS-related TIN and GMN

| Items | Total (n=39) | TIN (n=21) | GMN (n=18) | P |
| --- | --- | --- | --- | --- |
| Age at KB (years, M±SD) | 42±14.1 | 36±12.5 | 48±13.5 | 0.008 |
| Male:female ratio | 9/30 | 3/18 | 6/12 | NS |
| PSS duration (years, IQR) | 5.0 [2.0, 16.0] | 3.0 [1.75, 7.50] | 11.5 [2.0, 20.0] | 0.025 |
| IS therapy (%) | 15/39 (38.5) | 8/21 | 7/18 | NS |
| ANA (+) (%) | 37/39 (94.9) | 21/21 | 16/18 | NS |
| ANA 1:1280 (+) (%) | 8/39 (20.5) | 5/21 | 3/18 | NS |
| ANA 1:640 (+) (%) | 16/39 (41.0) | 11/21 | 5/18 | NS |
| Anti-SSA (+) (%) | 34/39 (87.2) | 18/21 | 16/18 | NS |
| Anti-SSB (+) (%) | 13/39 (33.3) | 9/21 | 4/18 | NS |
| Serum IgG elevation (%) | 25/38 (65.8) | 17/20 | 8/18 | 0.016 |
| Serum IgA elevation (%) | 14/38 (36.8) | 6/20 | 8/18 | NS |
| Serum IgM elevation (%) | 4/38 (10.5) | 3/20 | 1/18 | NS |
| Hypocomplementemia |  |  |  |  |
| Low CH50 | 3/38 (7.9) | 0/20 | 3/18 | NS |
| Low C3 | 6/38 (15.8) | 1/20 | 5/18 | NS |
| Low C4 | 6/35 (17.1) | 2/17 | 4/18 | NS |
| RTA(%) | 22/39 (56.4) | 20/21 | 2/18 | <0.001 |
| Urinary calcification/stone (%) | 12/39 (30.8) | 10/21 | 2/18 | 0.018 |
| Microhaematuria (%) | 18/39 (46.2) | 5/21 | 13/18 | 0.004 |
| Proteinuria (%) |  |  |  |  |
| Mild (<1.5 g/day) | 25/39 (64.1) | 17/21 (81.0) | 8/18 (44.4) |  |
| Moderate (1.5-3.5 g/day) | 7/39 (17.9) | 4/21 (19.0) | 3/18 (16.7) | 0.006 |
| Nephrotic range (>3.5 g/day) | 7/39 (17.9) | 0/21 (0) | 7/18 (38.9) |  |
| Urine collection (g/day, M±SD) | 2.2±3.7 | 0.8±0.6 | 3.9±4.9 | 0.033 |
| Scr (μmol/L, M±SD) | 103.9±40.99 | 113.2±42.33 | 93.0±37.60 | NS |
| eGFR(mL/min.1.73m<sup>2</sup>, M±SD) | 70.2±27.23 | 64.0±28.17 | 77.5±24.90 | NS |

NS, not significant. TIN, tubular interstitial nephritis. GMN, glomerulonephritis. KB, kidney biopsy. IS, immunosuppression. ANA, anti-nuclear antibodies. RTA, renal tubular acidosis. Scr, serum creatinine. eGFR, estimated glomerular filtration rate (calculated with the CKD-EPI creatinine equation). M±SD, mean±standard deviation. IQR, interquartile range.

Compared with patients with GMN, patients in the TIN group were younger and exhibited a shorter duration of pSS (TIN 3.0 [1.75, 7.50] years vs. GMN 11.5 [2.0, 20.0] years, P=0.025), and more frequently exhibited elevated IgG levels (TIN 17/20 vs. GMN 8/18, P=0.016). Patients with GMN exhibited more prominent microscopic haematuria (TIN 5/21 vs GMN 13/18, P=0.004) and proteinuria (TIN 0.8±0.6g/day vs. GMN 3.9±4.9g/day, P=0.033). Renal function at kidney biopsy did not differ between the TIN and GMN groups (Table 1).

### 2. Kidney biopsy findings and glomerular C4d deposition

Among the 18 patients with GMN, 8 were diagnosed with MN, 5 with MePGN, 2 with IgAN and 3 with MCD. Twenty-one patients were diagnosed with TIN. Specifically, 2 patients were diagnosed with acute interstitial nephritis, and the other 19 were diagnosed with chronic interstitial nephritis.

All 8 patients with MN exhibited C4d deposition along the glomerular capillary and/ or in mesangium (Fig 2 A, B). IF staining revealed IgG deposition in 7/8 of patients with MN and C3 deposition in 2/8 of patients with MN. Only 1 patient displayed weak C1q deposition using IF staining. Sub-epithelial electron-dense deposits (EDDs) and occasional mesangial EDDs were also observed in patients with MN using EM, including the patient with negative IgG deposition, confirming the diagnosis of MN (Table 2). Six of the patients with MN were diagnosed with stage II-III disease and exhibited different degrees of mesangial proliferation. Four cases had segmental thickening of GBM. Three out of five patients with MePGN and 1/2 patients with IgAN had mild mesangial deposition of C4d, but the staining area did not meet the criteria for glomerular C4d positive (Fig 2 C).

**Table 2:**
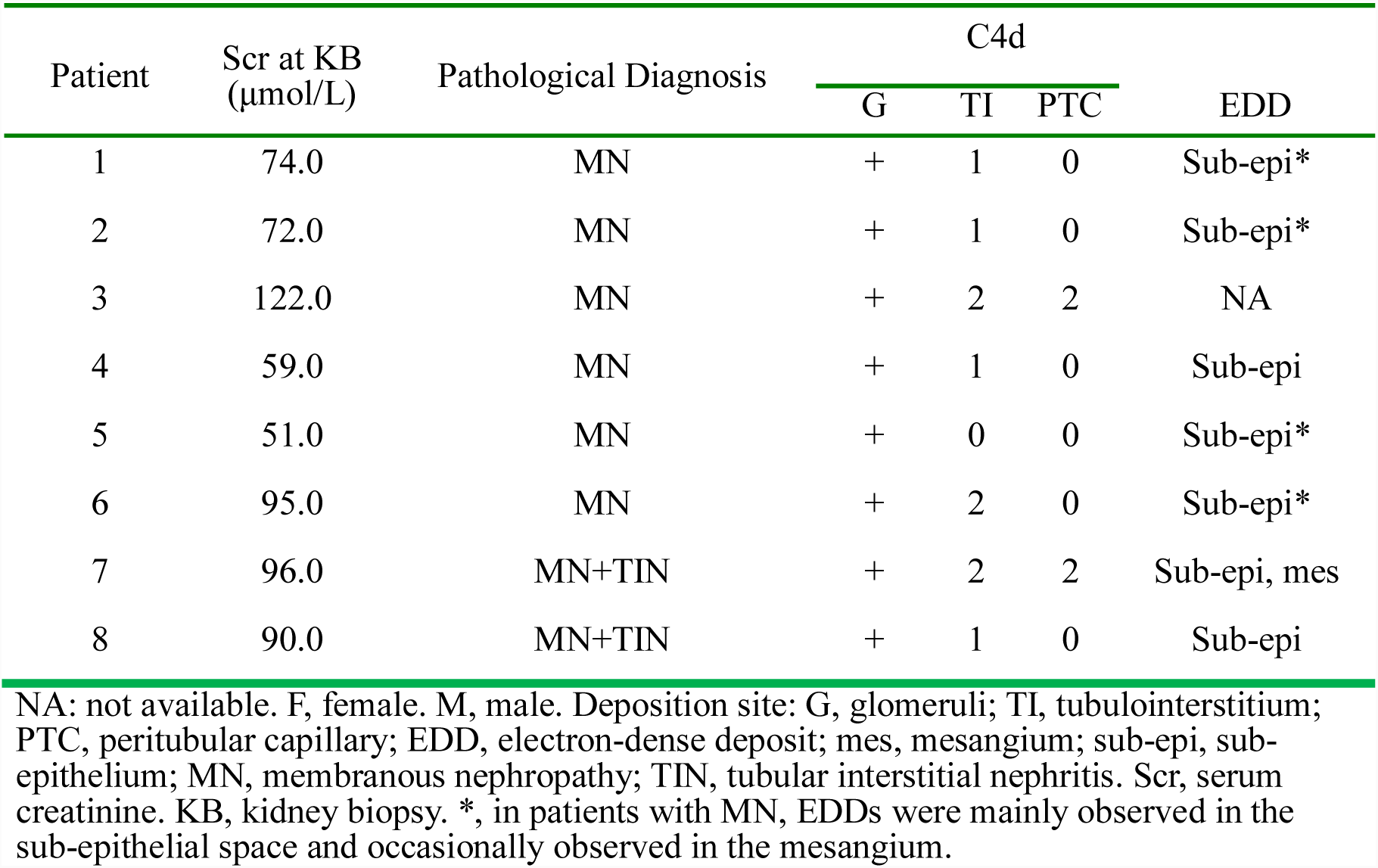
Details of C4d deposition in patients with pSS-MN

**Figure 2.**
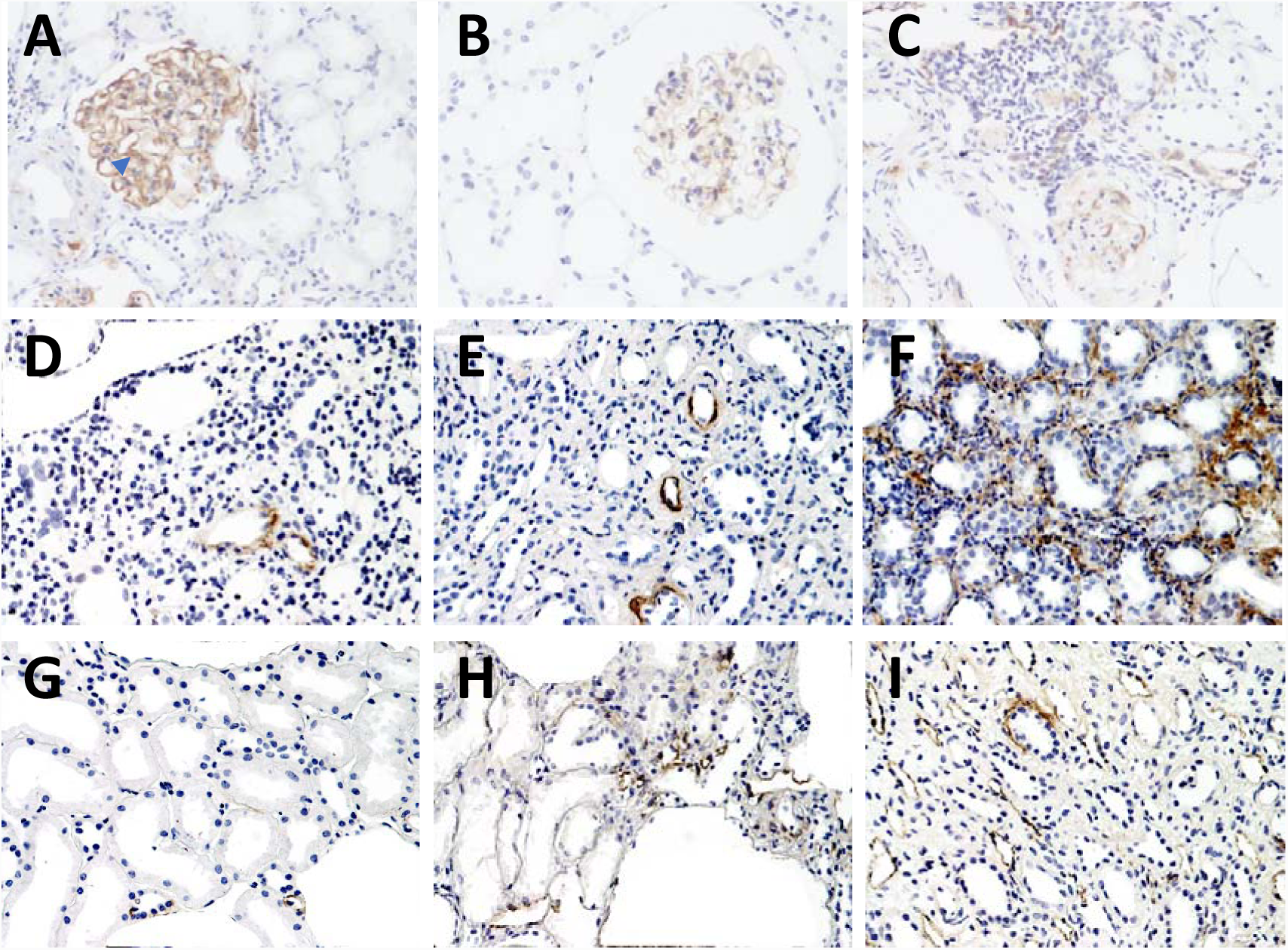
C4d deposition in kidney tissues from patients with pSS related renal impairments pSS renal impairments (light microscopy, IHC). Tubulointerstitial C4d and peritubular capillary C4d staining were semi-quantitatively scored from 1 to 3. A×200, pSS-related MN, C4d continuous staining along GBM and mild staining in mesangium(arrow). B×200, pSS-related MN, C4d segmental staining along GBM. C×100, pSS-related IgAN, mild mesangial staining of C4d which did not meet the criteria for G-C4d+, mesangial deposition of IgA and C3 positive. D×200, TI-C4d score 1, clinically dRTA, TIN, IF(-). E×200, TI-C4d score 2, clinically dRTA, TIN, IF(-). F×200, TI-C4d score 3, clinically RTA, TIN+mild MePGN, glomerular IF(-), interstitial C3 deposition positive by IF. G×200, PTC C4d score 1(minimal staining), TIN+MePGN, IF(-). H×200, PTC C4d score 2(focal staining), early MN, with IgG(3+), IgA(2+) and C3(3+) deposition along GBM, C1q negative. I×200, PTC-C4d score 3(diffuse staining), TIN.

### 3. Tubulointerstitial and peritubular capillary C4d deposition and clinical and pathological differences between groups

Tubulointerstitial C4d deposition was observed in 32 patients (17/21 patients with TIN and 15/18 patients with GMN), including 16 patients with weak or spotty staining, 12 patients with patchy C4d staining, and 4 patients with diffuse C4d staining (Fig 2 D, E and F). Two of the 4 patients displaying diffuse tubulointerstitial C4d deposition exhibited tubulointerstitial IgG or C3 deposition. Immune complex deposition was relatively rare in patients with TIN. Four patients with TIN exhibited tubular and interstitial deposition of IgG and C3.

We compared the clinical and pathological characteristics between the tubulointerstitial C4d-positive group (TI-C4d score >1) and C4d-negative group (Table 3). TI-C4d-positive patients experienced a longer duration of renal involvement (P=0.001) and exhibited increased serum creatinine levels at kidney biopsy (P=0.008). TI-C4d score had a linear correlation with serum creatinine levels at kidney biopsy (β=17.5±6.92, P=0.016). Pathologically, the TI-C4d^+^ group exhibited a higher interstitial fibrosis score (P=0.035) (Table 3). The TI-C4d score exhibited weak correlations with the percentage of sclerotic glomeruli (Spearman’s Rho=0.432, P=0.013), the degree of interstitial infiltration of mononuclear cells (Rho=0.339, P=0.035), and the degree of interstitial fibrosis (Rho=0.380, P=0.017).

**Table 3:** Comparison between the C4d-positive and C4d-negative groups

| Items | TI-C4d<sup>+</sup> (n=16) | TI-C4d<sup>-</sup> (n=23) | P |
| --- | --- | --- | --- |
| Age (years, M±SD) | 41±12.6 | 42±15.4 | NS |
| SS duration (years, M±SD) | 9.9±9.17 | 8.4±9.91 | NS |
| Renal duration (years, IQR) | 4.0 [2.0, 9.3] | 0.9 [0.1, 2.0] | 0.001 |
| ANA positive | 16/16 | 21/23 | NS |
| ANA 1:1280 | 5/16 | 3/23 | NS |
| Anti-SSA (+) | 13/16 | 21/23 | NS |
| Anti-SSB (+) | 6/16 | 7/23 | NS |
| Serum IgG elevation | 11/15 | 14/23 | NS |
| Serum IgA elevation | 8/15 | 6/23 | NS |
| Serum IgM elevation | 3/15 | 1/23 | NS |
| Hypocomplementemia Low CH50 | 2/16 | 1/22 | NS |
| Low C3 | 4/16 | 2/22 | NS |
| Low C4 | 3/15 | 3/20 | NS |
| CRP elevation | 3/13 | 0/19 | 0.058 |
| Scr (μmol/L, IQR) | 132.5 [89.7, 165.5] | 83.0 [70.7, 102.0] | 0.008 |
| eGFR (mL/min.1.73 m<sup>2</sup>, M±SD) | 58.7±27.31 | 78.3±24.65 | 0.025 |
| 24-h urine collection (g/day, M±SD) | 1.9±2.09 | 2.5±4.48 | NS |
| Glomerular sclerosis (% , IQR) | 19.1 [0, 49.6] | 4.2 [1.9, 6.3] | 0.051 |
| Interstitial fibrosis (M±SD) | 2.3±1.29 | 1.4±1.04 | 0.035 |

NS, not significant. TI-C4d, tubulointerstitial C4d. SS, Sjögren's syndrome. ANA, anti- nuclear antibodies. CRP, C reactive protein. Scr, serum creatinine. eGFR, estimated glomerular filtration rate (calculated with CKD-EPI creatinine equation). M±SD, mean±standard deviation. IQR, interquartile range. Tubular atrophy and interstitial fibrosis were scored according to the percentage of interstitial area involved: 0 (absent), 1 (≤25%), 2 (25-50%), 3 (50%-75%), and 4 (>75%).

PTC C4d deposition was observed in 12 patients, including 4 patients with minimal deposition (Fig 2 G), 4 patients with local deposition (Fig 2 H) and 4 patients with diffuse deposition (Fig 2 I). The PTC C4d score was positively correlated with the ANA titre (Spearman’s Rho=0.458, P=0.003). All of 4 patients with diffuse deposition had elevation of serum IgG levels, and 3 of them had concomitant elevation of serum IgA levels and ANA titre at 1:1280, the serum levels of IgM were within normal range for all 4 patients.

### 4. Treatment and patient follow-up

Thirty-seven patients received glucocorticoids. Twenty-one patients simultaneously received cyclophosphamide therapy. Other immunosuppressive agents included cyclosporine A(n=1), mycophenolate mofetil(n=1), azathioprine(n=1), methotrexate(n=6), leflunomide(n=1), hydroxychloroquine(n=1) and tripterygium glycosides (n=7). The median follow-up time was 642 [239-1458] days. Ten (25.6%) patients exhibited a 20% increase in eGFR at the end of follow-up. The 4 patients with PTC C4d diffuse deposition were treated with prednisone and immunosuppressants (leflunomide in one patient, methotrexate in another and cyclophosphamide in the other two). Two of them were followed for more than half a year and did not exhibit a significant improvement of renal function. In one patient, Scr at the time of kidney biopsy and of the last follow-up time were 164μmol/L (eGFR 44mL/min.1.73 m^2^) and 177μmol/L (eGFR 40mL/min.1.73 m^2^), respectively. In the other patient, Scr varied from 97μmol/L (eGFR 57mL/min.1.73 m^2^) at the time of kidney biopsy to 86μmol/L (eGFR 66mL/min.1.73 m^2^) at the end of follow-up.

## Discussion

This study investigated the clinical significance of renal C4d deposition in patients with pSS-related renal lesions which has not been reported before. Patients with pSS-MN exhibited glomerular C4d deposition without obvious concomitant C1q deposition, indicating the involvement of the lectin pathway of complement activation in pSS-MN. Tubulointerstitial and PTC C4d deposition were also observed. The tubulointerstitial-C4d score was correlated with kidney function at the time of kidney biopsy, indicating a possible link between renal interstitial injury and autoantibody-mediated complement activation in patients with pSS.

GMN was more prevalent in our study (46.2%) than in previous studies using serum and urine biochemical examinations (13.9-29%)(Bossini et al., 2001;Ren et al., 2008). In studies employing pathological investigations, the proportion of GMN ranged from 37.1 to 48.6%(Goules et al., 2013;Kaufman et al., 2008;Kidder et al., 2015). In the GMN group, there were 8 patients with the pathological diagnosis of MN. The PLA2R/THSD7A antibodies in combination with IgG4 is a useful tool to diagnose intrinsic antibodies related MN(L’Imperio et al., 2018).As the retrospective nature of our study, there were not enough biopsy tissues for us to test the expression of PLA2R/THSD7A antibodies as well as IgG subtypes. However, 6 of the 8 patients with MN presented with mesangial proliferation and occasional mesangial EDDs were observed in 5 patients, 4 of them had segmental thickening of GBM, which are more likely features of secondary MN(Larsen et al., 2013). One patient with negative IF was confirmed by detecting sub-epithelial EDDs under the EM. The IF negativity was possibly due to the early disease stage and lack of abundant immune complex. No obvious clinical and pathological features of lupus nephritis were observed both at the time of kidney biopsy and during follow up, we excluded the possibility of MN secondary to SLE. MN was the most common type of glomerular involvement in our study. In previous studies, MPGN was the most common type of GMN(Kidder et al., 2015;Matignon et al., 2009) and was typically accompanied by cryoglobulinemia. The reason for the increased proportion of patients with MN in our study may be due to differences in environmental and genetic factors, because the Chinese population exhibits an increased prevalence of MN and less prevalence of MPGN(Pan et al., 2013).

The contribution of complement to MN (primary and secondary) was documented elsewhere(Ma, Sandor &Beck, 2013). Besides C3, the component of the alternative pathway, glomerular deposition of C4d and MBL were detected in PMN, indicating involvement of the lectin pathway(Hayashi et al., 2018;Hui et al., 2014). The lectin pathway of complement may be possibly triggered by the in situ formed immune complexes comprising the IgG4 type of anti-PLA2R antibodies(Sinico et al., 2016). In patients with SLE-MN, circulating immune complexes deposition and the subsequent activation of the complement system resulted in the full house deposition of immune complexes and C1q, and all the three pathways of complement activation were involved, namely the classical, lectin(Kim et al., 2013) and alternative pathway(Song et al., 2017). All patients with MN in our study exhibited glomerular C4d deposition without obvious C1q deposition, indicating local complement activation by the lectin pathway, which is similar to PMN(Espinosa-Hernández et al., 2012) and different from SLE-MN(Kim &Jeong, 2003), the underlying mechanism remains to be investigated.

The significance of tubulointerstitial C4d deposition is not clear, and few studies are available. Tubular C4d deposition was associated with an increased grade of WHO classification of IgAN(Maeng et al., 2013). Patients with TBM C4d deposition exhibited a significantly higher score for the viral cytopathic effect on BK nephropathy(Batal et al., 2009). In patients with lupus nephritis, no practical significance of TBM-C4d was identified(Batal et al., 2012). We identified spread C4d deposition along TBM and in the renal interstitial compartment with different degree. The semi-quantified TI-C4d score was correlated with worse renal function at kidney biopsy and more severe interstitial fibrosis, suggesting potential contribution of complement activation to tubulointerstitial injuries and renal impairment(Abbate et al., 2008). For most patients with pSS-related TIN, the IF staining results were negative and cellular immunity is considered a main pathological mechanism. The occasional TBM deposition of complement in images of IF staining is typically considered nonspecific(Evans et al., 2016). Technically, immunohistochemistry for TBM C4d in paraffin-embedded renal allograft tissues is more specific than IF in frozen tissues (4% by IHC vs. 48% by IF)(Batal et al., 2008). Thus, we considered that the tubulointerstitial C4d deposits identified by IHC in our study were more inclining to suggest its potential role in the pathoetiological aspect. Due to the complexity of the complement system in diseases(Ricklin, Reis &Lambris, 2016), the cause-effect relationship of complement activation and renal lesions in patients with pSS can not be demonstrated in this study, which certainly is an area needing to be further investigated.

As a marker for antibody-mediated renal allograft rejection(Regele et al., 2002), PTC C4d has been less frequently reported in glomerulonephritis, and different pathological mechanisms may be involved. In patients with LN, local immune complexes may be responsible for PTC C4d deposition as EDDs along the basement membrane of PTC were identified in 24/31 (77.4%) of patients with LN presenting PTC C4d deposition, whereas none of the patients with acute antibody-mediated rejection exhibited deposition(Li et al., 2007). In addition to autoantibodies, local shear stress on endothelial cells may play a role in the renal crisis in patients with scleroderma(Okroj et al., 2016;Yin et al., 2008), and PTC C4d correlated with an increased risk of renal failure(Batal et al., 2009). In our study, PTC C4d deposition was identified in 12 patients. The PTC C4d score was positively correlated with the ANA titre. Thus, PTC C4d deposition in pSS-related lesions was possibly mediated by the effects of autoantibodies, which is in accordance with the detection of anti-carbonic anhydrase II antibodies(Takemoto et al., 2005) and antibodies against intercalated cells(Devuyst et al., 2009) in pSS-related renal tubular lesions. As few patients exhibited diffuse PTC C4d deposition, the significance of PTC C4d deposits in pSS-related renal injury remains to be further clarified.

In consistence with Kidder et al’ s study showing similar renal survival rate between the GMN and the TIN pathological entities(Kidder et al., 2015), our study also showed that renal function at kidney biopsy in patients with GMN did not differ from those with TIN. PTC C4d deposition was a prognostic marker in renal allograft(David-Neto et al., 2007). Although 9 patients in our study exhibited improvement of renal function, two of the 4 patients with PTC C4d diffuse deposition did not show significant improvement of renal function during follow-up, indicating poor response to treatment. The other two patients with PTC C4d diffuse deposition were lost follow up shortly after discharge, which resulted in the small sample size, the impact of C4d on renal function in patients with pSS related renal impairments remains to be explored in a prospective study with large sample size.

The retrospective nature of this study may lead to possible bias, as more patients with GMN were listed in the kidney biopsy database. Other limitations of our study were that we did not examine the serum anti-PLA2R antibody titre, and an IgG sub-type analysis was not performed in renal tissues. The inadequacy of biopsy tissues hindered the testing of other complement components (such as lectin and C5b-9).

## Conclusions

This preliminary study investigated C4d deposition in patients with pSS with renal involvement. Glomerular C4d positivity was detected in 100% of patients with pSS-MN, in the absence of C1q, indicating possible complement activation through the lectin pathway. The tubulointerstitial C4d deposition observed in our patients with pSS indicated possible involvement of the MBL pathway of complement activation in pSS-renal lesions. Tubulointerstitial C4d deposits might be a candidate of pathological markers of severe renal injury in patients with pSS-related renal impairments, which remains to be further investigated.

## Supporting information

Positive control

Negative control

## Acknowledgements

The authors thank Dr. Jianling Tao (Department of Medicine, Stanford University) for editing the manuscript.

