## Supplementary figures and images for "Clinical significance of C4d deposition in renal tissues from patients with primary Sjögren’s syndrome—A preliminary study"

### Negative control

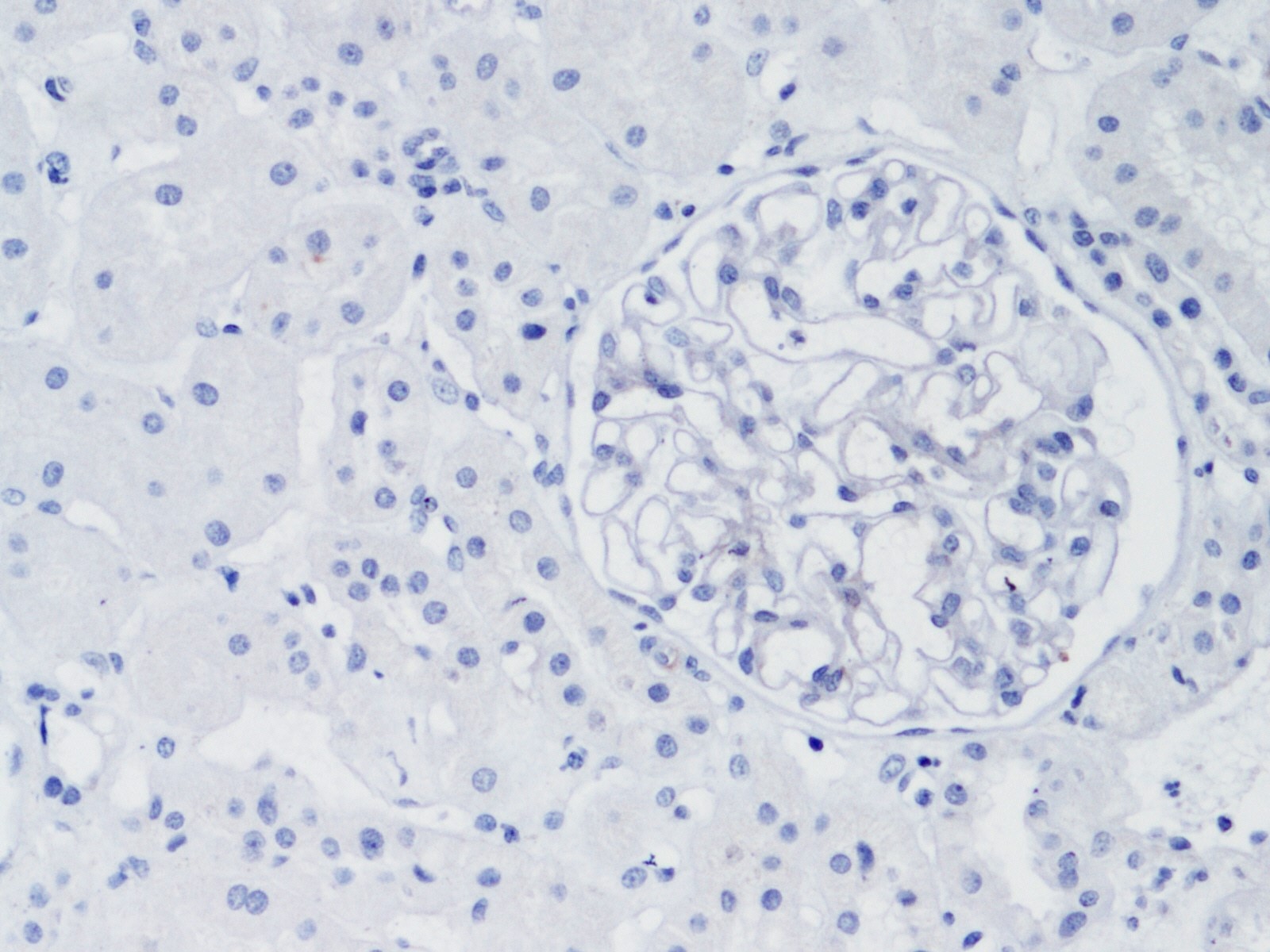

### Positive control

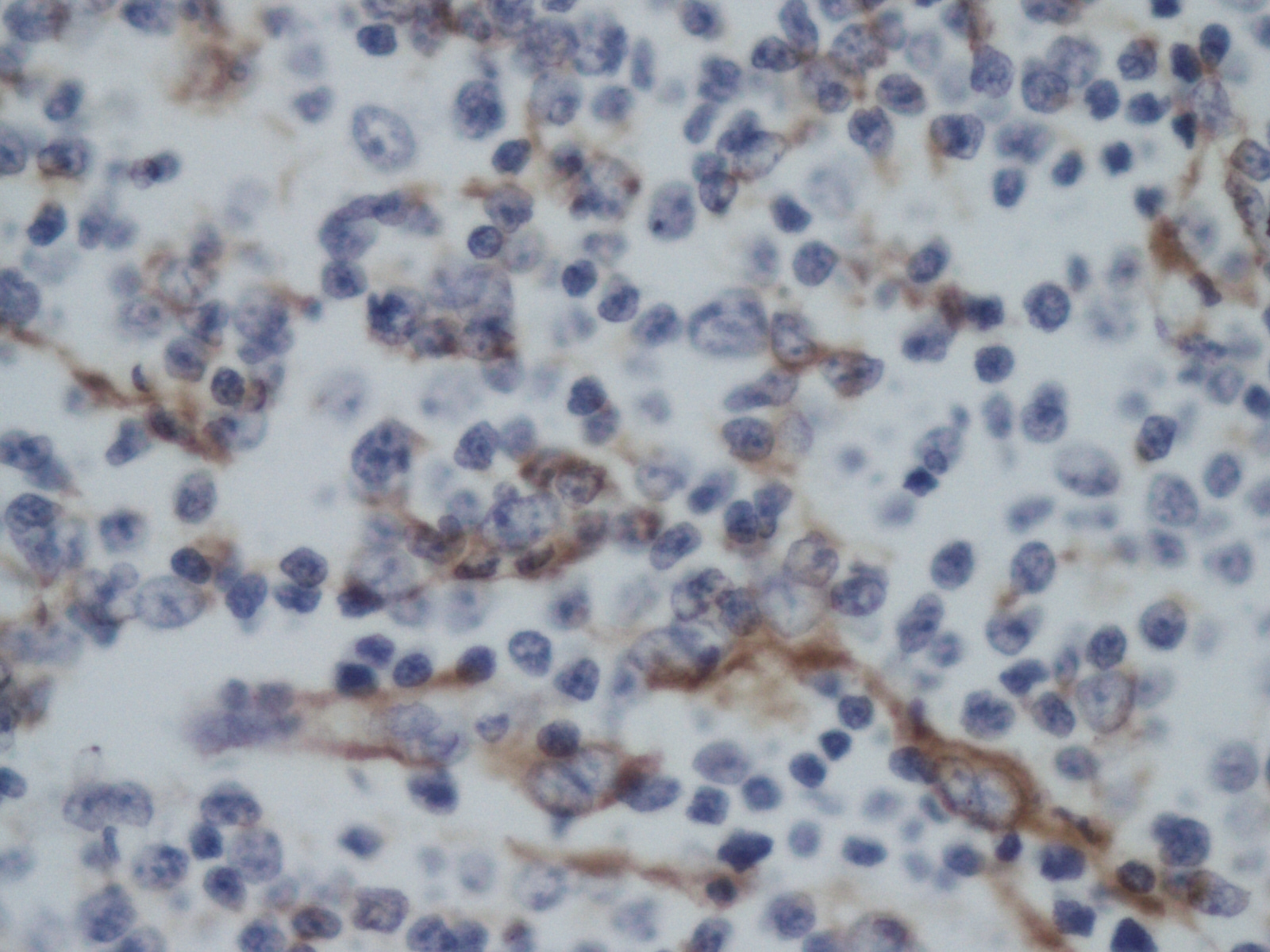
